# Do perineuronal nets stabilize the engram of a synaptic circuit?

**DOI:** 10.1101/2023.04.09.536164

**Authors:** Varda Lev-Ram, Sakina P. Lemieux, Thomas J. Deerinck, Eric A. Bushong, Brandon H. Toyama, Alex Perez, Denise R. Pritchard, Sung Kyu R. Park, Daniel B. McClatchy, Jeffrey N. Savas, Susan S. Taylor, Mark H. Ellisman, John Yates, Roger Y. Tsien

## Abstract

Perineuronal nets (PNN), a specialized form of ECM (?), surround numerous neurons in the CNS and allow synaptic connectivity through holes in its structure. We hypothesis that PNNs serve as gatekeepers that guard and protect synaptic territory, and thus may stabilize an engram circuit. We present high-resolution, and 3D EM images of PNN- engulfed neurons showing that synapses occupy the PNN holes, and that invasion of other cellular components are rare. PNN constituents are long-lived and can be eroded faster in an enriched environment, while synaptic proteins have high turnover rate. Preventing PNN erosion by using pharmacological inhibition of PNN-modifying proteases or MMP9 knockout mice allowed normal fear memory acquisition but diminished remote-memory stabilization, supporting the above hypothesis.

**Significance:** In this multidisciplinary work, we challenge the hypothesis that the pattern of holes in the perineuronal nets (PNN) hold the code for very-long-term memories. The scope of this work might lead us closer to the understanding of how we can vividly remember events from childhood to death bed. We postulate that the PNN holes hold the code for the engram. To test this hypothesis, we used three independent experimental strategies; high-resolution 3D electron microscopy, Stable Isotop Labeling in Mammals (SILAM) for proteins longevity, and pharmacologically and genetically interruption of memory consolidation in fear conditioning experiments. All of these experimental results did not dispute the PNN hypothesis.

## Introduction

The molecular and cellular basis for very-long-term memory is among the most central, captivating, and controversial questions in neuroscience. It requires both intra- and extracellular changes that contribute to memory persistence in the brain over time(1, 2). Richard Semon coined the term “*engram”* and in his theory, these memories leave a physical mark comparable to engraved writing in the brain (physical engraving-see below) (3). We hypothesize that the PNN, a specialized extracellular matrix (ECM), is the substance that undergoes this physical engraving to stabilize the *engram*. The PNN are modified during synapse strengthening, a process important for synapse stability and memory. Studies that rely on removing chondroitin (a major component of PNN/ECM structure) by injecting the enzyme chondroitinase ABC (ChABC) into the brain suggest that PNN/ECM is essential for preserving remote-memories(1, 4–6). The schematic model in figure 1 illustrates elements of PNN that may be altered (“edited”)following synaptic activation during memory consolidation or long-lasting memory stabilization. Simply stated, learning-linked synaptic changes involve release of matrix metalloproteases (MMPs) in the vicinity of the activated synapses, breaking up and eroding adjacent PNN(7).

**Figure 1.**
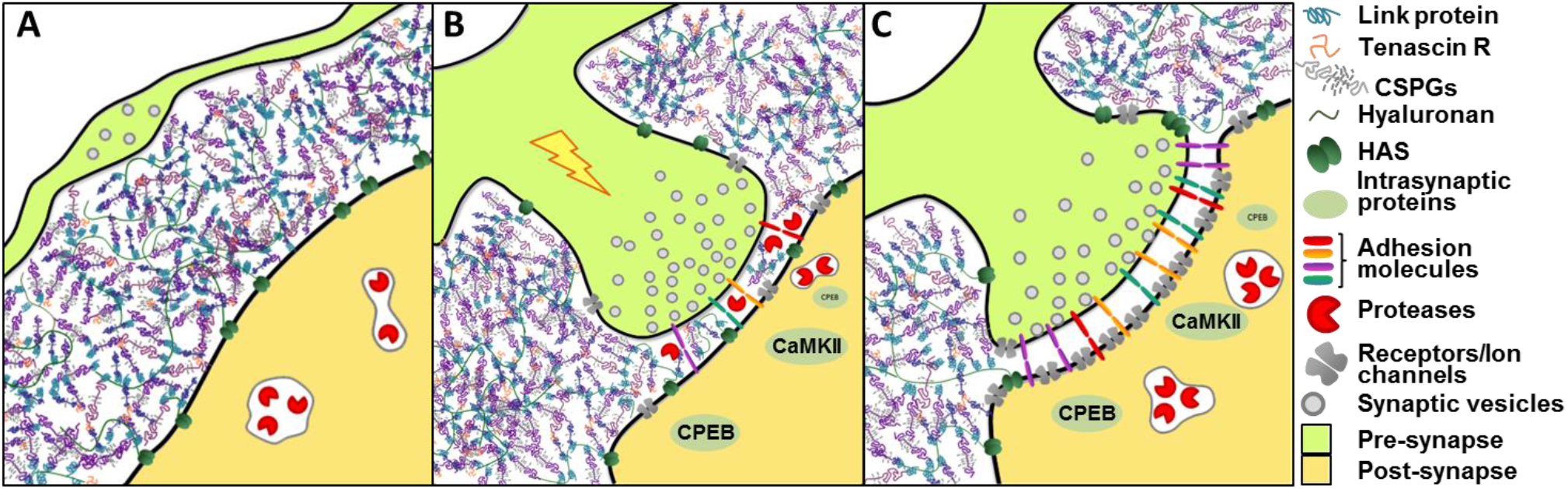
Schematic model of PNN. (**A**) PNN creating a physical barrier that limits synaptogenesis in adult animals. (**B**) Learning-associated synaptic activity leads existing synapses to release matrix metalloproteases from the post-synaptic cell facilitating synaptic territory enlargement and stabilization. (**C**) Stable and strong synapse with a larger hole in the PNN the might support a stronger and long-lasting connection.

Roger Tsien suggested that this process might be very local and could result in more permanent “real-estate” for the activated synapse thereby, stabilizing and protecting the territory of the larger and stronger synapse(8). He speculated that the code for a memory might reside in the pattern of holes etched in the stable PNN coating, analogous to old computer punch cards where holes permit contact between brushes placed on both sides of the card that allows an electric current flow, like a synaptic connection. Unlike the computer punch cards, the size of the hole in the PNN indicates the allowance of synaptic connection and hence the strength of the synapse. This model would require that the stability and longevity of the PNN component is much greater than synaptic proteins.

By using SILAM, we determined that the degradation dynamics of PNN proteins resembles some of the known long-lasting proteins (figure 3)(9). We also labeled the PNN for imaging with antibodies for aggrecan, or with Wisteria Floribunda Agglutinin (WFA). WFA has a high affinity to N-acetylgalactosamines (GalNAc) branches attached to the glycosaminoglycans(GAGs)(10),WFA staining revealed a thick, fenestrated structure around some neurons in fixed adult brain tissue. This stain specifically labels material surrounding fast-spiking parvalbumin-rich inhibitory interneurons(11, 12), and a subset of excitatory neurons, such as pyramidal cells in the hippocampal CA2 area(13) and in the basolateral amygdala(14). Here, we used structural, proteomic, and behavioral approaches to examine this hypothesis.

## Results

### Observation by high-resolution EM and 3D-EM confirms that the PNN holes are occupied by synapses

To test the hypothesis that the pattern of PNN-holes holds the code for very-long-term memories, we first tested whether synapses occupy all or most of the PNN holes. The above observation has been reported using light microscopy (LM) in cultured neurons(15) and *in vivo*(14), but this has never been confirmed using high-resolution 3D-EM.

We applied advanced 3D-EM imaging technologies to examine the relationship between synapses and PNN holes(12). Using antibodies for aggrecan (Ab) in the deep cerebellar nucleus (Fig.2A1-3) and WFA-biotin in the hippocampal CA2 region (Fig.2B1-3), we heavily labeled PNN-engulfed neurons that exhibited openings that are only occupied by synapses (Fig. 2A3&2B3). To get a better understanding of the PNN-synapse relationship, we used serial block-face scanning electron microscopy (SBEM) to create high-resolution 3D-EM(16). We manually segmented the plasma membrane and semi-automatically marked the PNN of neurons and dendrites from a hippocampal CA2 volume. In figure 2C1, the dendrite (white) is surrounded by the porous PNN (magenta).

**Figure 2.**
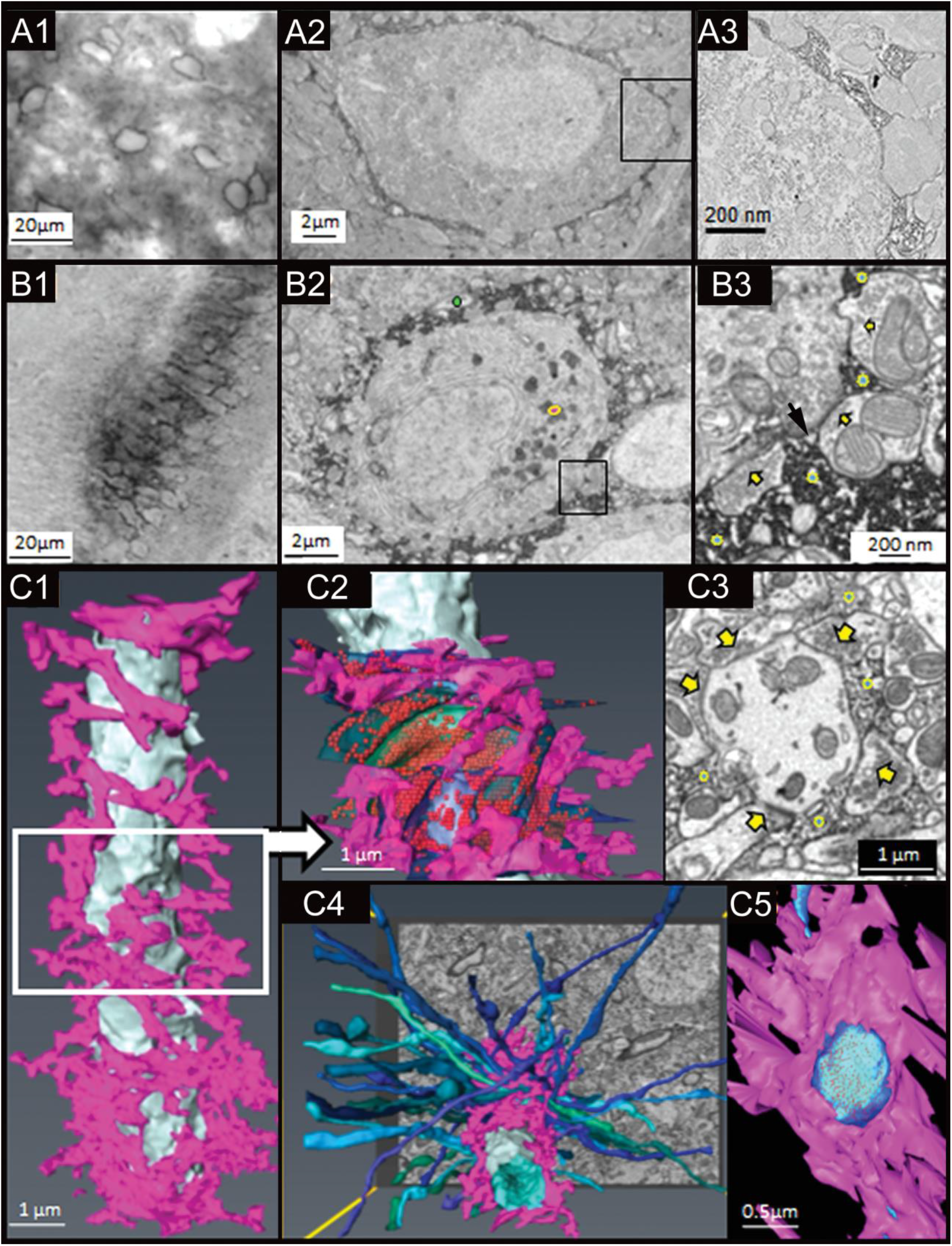
Holes in PNN allocate spaces for synaptic contact. (**A**) PNN in the deep cerebellar nucleus labeled with Ab for aggrecan followed by biotinylated Ab and streptavidin-HRP. (**A**1) LM of PNN engulfed neurons. (**A**2) EM of a PNN surrounded neuron. (**A**3) Higher magnification demonstrating synapses delimited by PNN. (**B**) Hippocampus CA2 area neurons labeled with WFA-biotin followed by streptavidin-HRP. (**B**1,2,3) are like **A**1,2,3, with symbols that demonstrate the different components. In **B**2,**B**3,&**C**3 PNN are labeled marked with (), synapses (), myelin (), and lipofuscin (). (**C**) Three dimensional reconstruction of a PNN engulfed dendrite segmented from a serial block face scanning electron microscope volume. (**C**1) Dendrite colored in white and PNN in fascia. (**C**2) Synaptic boutons are highlighted in green and blue, with red synaptic vesicles showing through the transparent plasma membrane. The segmentation highlights only the midsection of the dendrite volume. (**C**3) An EM image from the center of this area demonstrating the dendritic membrane, surrounded by either synapses or PNN. (**C**4) Top view of the dendrite with axons that synapse onto the mid-section. (**C**5) A view from the inside of the dendrite through a PNN hole demonstrating a single synapse contacting the dendrite.

We traced all the presynaptic boutons (translucent greens and blues) and neurotransmitter vesicles (red balls) within the center section of the dendrite. Interestingly, more than 98% of the dendrite plasma membrane was in contact with either PNN or presynaptic boutons. We did not observe contacts with other passing dendrites, myelinated fibers, or astrocytic processes (Fig.2C2). This is in contrast to "naked" dendrites without a PNN coating, that have membrane-to-membrane contact with myelinated fibers or other passing dendrites (Fig.S1).

The single-plane EM image in figure 2C3 demonstrates that the entire circumference of the dendrite is occupied by either PNN or synapses that in turn are restricted and protected by PNN. A three-dimensional view of the axons that synapse onto the PNN- engulfed dendrite is presented as a volume in figure 2C4. Figure 2C5 is a view from inside this dendrite through a hole in the PNN, exhibiting synaptic contact that is both confined and secured by the surrounding PNN. This supports the idea that to increase, and thus strengthen a PNN-surrounded synapse, the PNN needs enzymatic chiseling.

### PNN protein longevity

As some memories can last for an organism’s lifetime, we proposed that the substrate that holds the code for memory should be stable and have a slow turnover. Other hypotheses for the preservation of remote-memory postulate that key synaptic proteins including Calcium–calmodulin (CaM)-dependent protein kinase II (CaMKII)(2), protein kinase M-zita (PKMζ)(17), or cytoplasmic polyadenylation element binding protein (CPEB3)(18) store memory by passing the information to all subsequent generational copies of the protein aggregates within synaptic processes. For all the proteins that are suggested as memory mediators by these hypotheses, a faithful repetitive intergenerational copying would be necessary to maintain a steady state over time.

To test the differential stability of synaptic and PNN components in the brain, we designed a SILAM experiment(19), fully labeling mice proteins with ^15^Nitrogen (^15^N) for proteomic analysis. A schematic representation of the experimental design is detailed in figure 3A, and further described in the supplementary method section.

**Figure 3.**
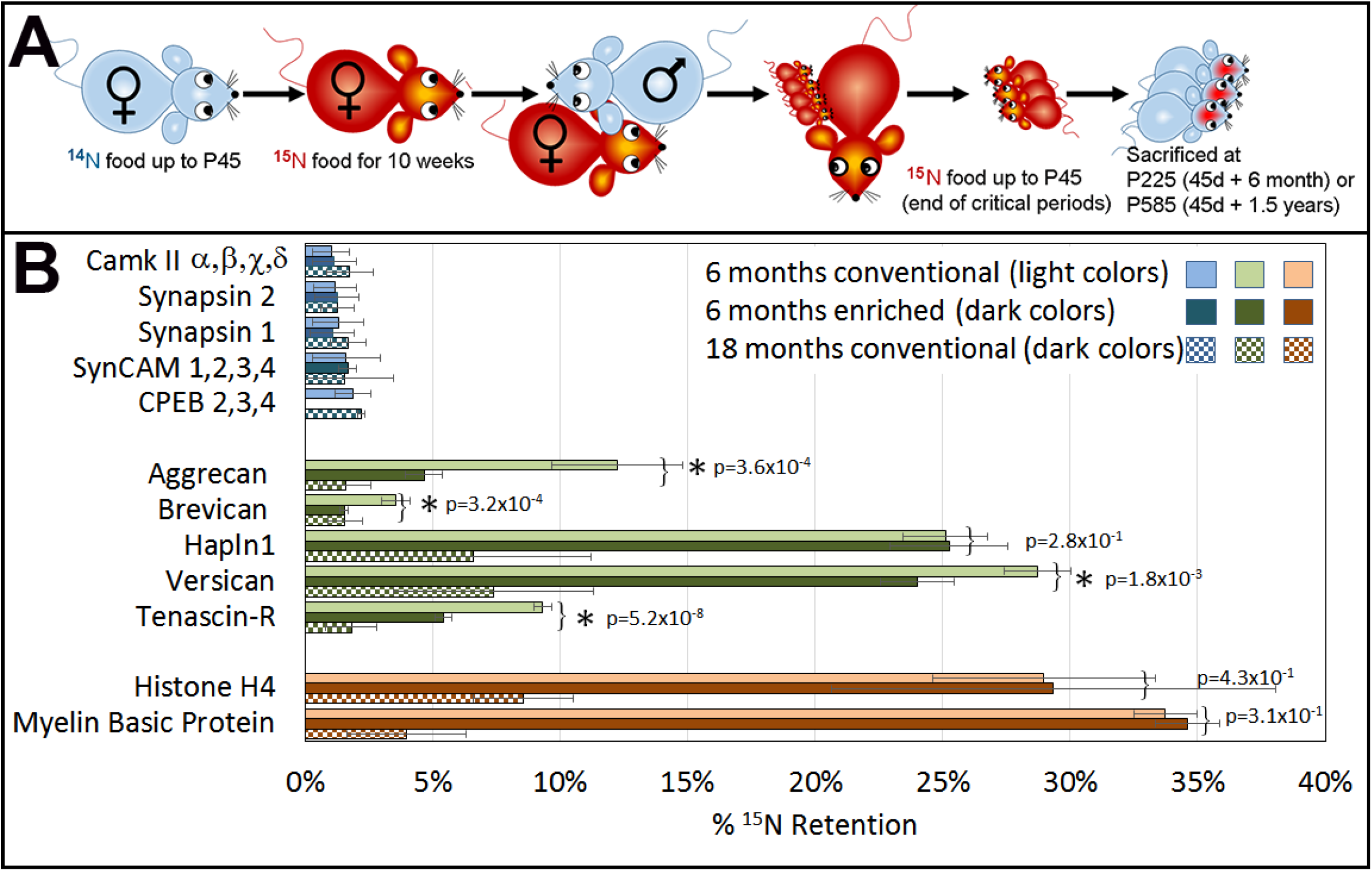
PNN proteins are long-lived but can be eroded by enriched environment as demonstrated by pulse-chase SILAM experiment. (**A**) Schematic representation of the experiment that is detailed in the supplementary material. (**B**) MuDPIT results demonstrating retention of ^15^N in stable proteins compared to proteins with fast turn-over rate. Light and dark color bars represent 6- and 18-months pulse-chase respectively. Histone-H4 and Myelin Basic Protein (MBP) are known stable proteins, and serve as a reference (orange). Synaptic proteins (blue) show low ^15^N fractional abundance even at the 6 months chase, while PNN proteins (green) retain a high level of ^15^N fractional abundance that is comparable to histone-H4 and MBP at 6 months pulse-chase. Some of the PNN proteins retained significant ^15^N fractional abundance even at the 18 months chase. Protein longevity comparison between EE and CC show synaptic proteins (checkered dark blues) did not demonstrate differences in turnover. PNN proteoglycans were eroded more in EE mice compared to littermates in CC, except for Hapln1 (checkered dark greens). While there was no change in known MBP and histone-H4 (checkered dark oranges). The number of peptides that were included in each group is detailed in Table S1.

The ^15^N pulse was chased by an ^14^N starting at P45. Mice were sacrificed at P45, P225, and P585 (0, 6, &18 months-^14^N chase). At each harvest point, we prepared synaptosomes and PNN enriched fractions and performed multi-dimensional protein identification technology (MudPIT) analysis to identify ^15^N retention by brain proteins.

Previous SILAM studies primarily focused on identification of intracellular proteins, while several extracellular proteins were also identified to be highly stable. The previous analysis was done in cultured neurons, and included; Tenascin-C, Tenascin-R, agrin, lamin(20, 21). In brain tissue the following long lasting molecules were identified;versican, Hapln1, and MBP (21, 22).

We analysed synaptic, PNN, and known long-lasting proteins. At the 6-months pulse-chase point, the synaptosomal proteins; CaMKII, synapsin, adhesion molecules (SynCAM), and CPEB; did not meet the criteria of ^15^N fractional abundance for long-lived proteins, as described in the material and methods section.

To achieve statistically meaningful data points, we averaged all isoforms of CaMKII, SynCAM, and CPEB. (Fig.3B, light blue bars). Meanwhile, constituents of the PNN; aggrecan, brevican, tenascin-R, versican, and Hapln1 retained 5%–28% ^15^N fractional abundance (Fig.3B light-green bars). We compare these data to proteins that are known to have a slow turnover like histone-H4 and myelin basic protein (MBP) that retained 28%-34% respectively (Fig.3B, light-orange bars). This is comparable to the 7–37% ^15^N that was previously found in other long-lasting structural proteins such as collagens and lamins(22).

In aged mice at 18 months pulse-chase, the PNN components were still clearly detectable at 2–7% ^15^N retention (Fig.3B, dark-green checkered bars). This was comparable to MBP and histone-H4, whose ^15^N had declined to 4% and 8% respectably (Fig.3B dark-orange bars). PNN components, MBP, and histone-H4, demonstrate a very slow turnover within a year, while synaptic proteins maintained the same levels of ^15^N retention as in the six months pulse-chase points (Fig.3B dark-blue checkered bars). This further shows that synaptic proteins values at both 6 and 18 months pulse-chase could not be considered as “heavy.” Hence, some of the PNN components are comparable to known long-lasting proteins in the brain whereas synaptic proteins have rapid turnover.

The six months pulse-chase group was divided into an enriched environment (EE, see bellow) and conventional cages (CC). We speculated that the mice in the EE; with changing toys, spinning disks, mirrors, balancing beams, tunnels, occasional music, scents, and frequent handling; would activate the brain and induce learning and memories that the CC group could not acquire. Thus, PNN erosion would result in less ^15^N retention in PNN proteins in EE caged mice. Indeed, we observed a statistically significant reduction in ^15^N retention in brains from EE mice (Fig.3C green bars) compared to the CC group (2-tailed t-test).

The known long-lived proteins that are represented here as a reference (histone-H4 and MBP) had the same turnover rate in EE and CC groups. This further validates the authenticity of the differences in PNN longevity between EE and CC. However, Hapln1s’ turnover was not accelerated due to EE, which might point to the preservation of the scaffolding of the hyaluronic acid net with Hapln1 tightly anchored.

### Remote-memory stabilization requires MMP(s) activity

PNN can be eroded by specific proteases that either create new holes for synaptogenesis or enlarge existing holes to expand the synaptic contact territory, and consequently strengthen synaptic input(23). The main proteases that are involved in this activity are MMP 2 and 9.

Fear conditioning has been demonstrated to evoke the expression of MMP-9 mRNA and enzyme activity in 3 major mouse brain structures; the hippocampus, prefrontal cortex, and amygdala(24). To test the role of MMPs in the establishment of long-lasting memories, we modified protease activity via either pharmacological inhibition or genetic knockout (KO). We performed classic fear conditioning, assessing the association of cues (tone and light) with an adverse mild electric shock.

Tests were performed 24 hours and four weeks after conditioning. We compared memory recall in four groups of mice wild-type (WT) and MMP9 KO, each group was injected with either a broad-spectrum MMP inhibitor, prinomastat (AG-3340), or with the vehicle (DMSO). Prinomastat has been demonstrated to cross the blood-brain barrier(25), and to inhibit MMP-2, -3, -9, -13, and -14(26). We also tested the presence of prinomastat in brains of injected animals. Tests were performed at 0, 2, 4, 8, and 24 hours after injection of 4mg prinomastat IP, using liquid chromatography mass spectroscopy x2 (LCMS2). We found up to 20nM prinomastat concentration of in the brain extract, 4 hours post injection (data not shown).

A Cue test performed at 24h post induction indicated that fear conditioning was equally acquired in all the groups. Nevertheless, testing the mice four weeks later revealed a large statistically significant difference of memory retention between the FVB WT that were injected with DMSO vs. prinomastat (2-tailed t-test p < 10^-4^; Fig.4B2). We concluded that MMPs inhibition might have prevented the long-lasting memory, more likely at the consolidation or stabilization stage rather than at the recall.

The FVB MMP9-KO mice had a significant deficiency in memory at the four weeks test, although the memory deficiency was lesser than the prinomastat injected WT. However, the KO mice memory at the 4 weeks test, did not differ between the DMSO and the prinomastat-injected groups. The partial memory in MMP9-KO mice in the four weeks test might be due to life-long compensation with prinomastat-insensitive proteases, because of the lack of MMP9 in their system (Fig.4B2).

We also tested the dynamics of remote memory in prinomastat-injected mice. Separate groups of mice were tested 1, 2, 3, 4, and 5 weeks after fear induction. We found decreased memory, which seemed to plateau at 4 weeks after induction (Fig 4B3). This might be due to fading recent memory, while the consolidation/stabilization of remote memory was impaired by MMP inhibition. Control groups injected with DMSO exhibited normal remote memory (Fig.S3).

**Figure 4.**
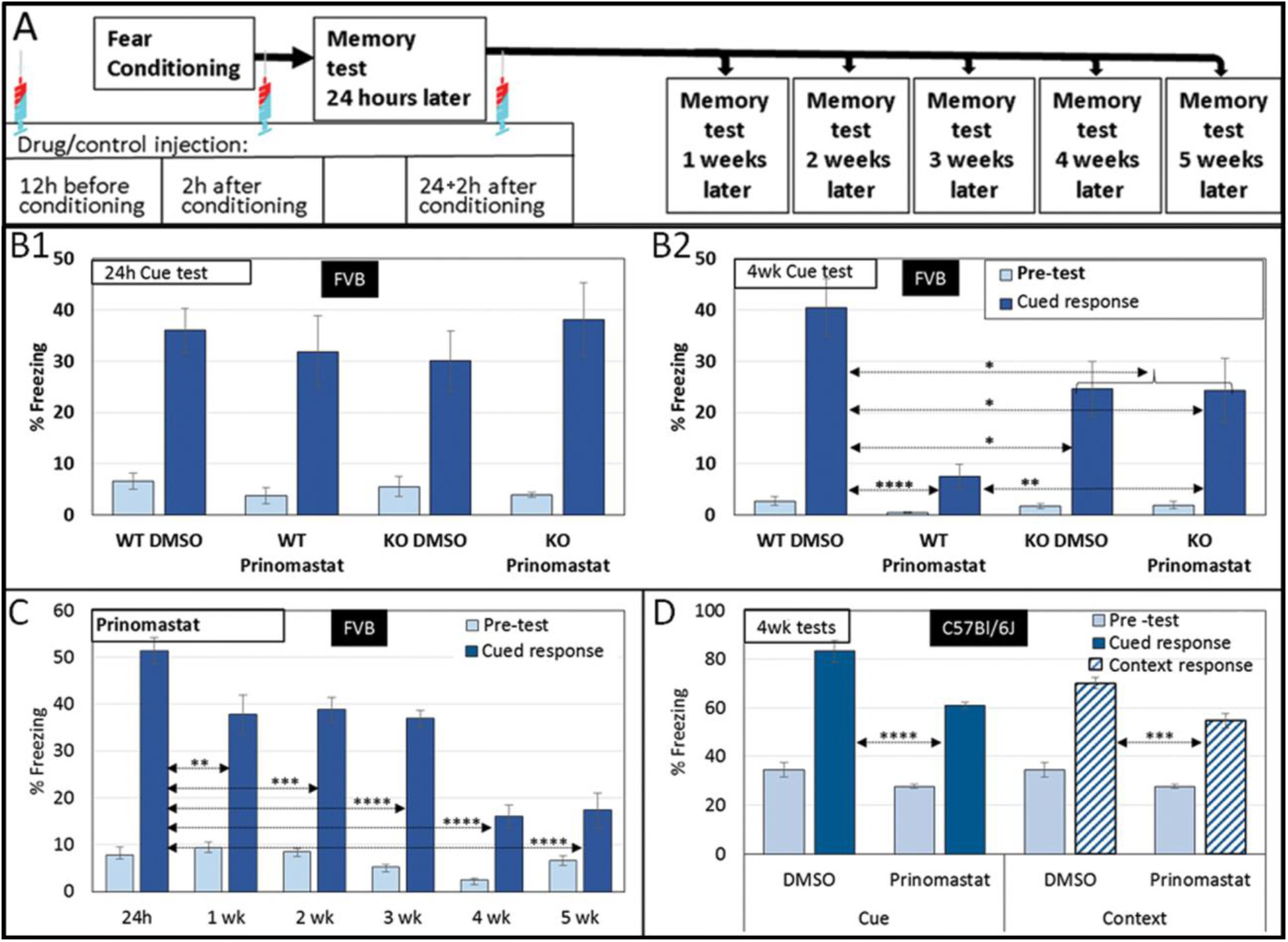
MMP inhibition effects remote-memory but not memory acquisition. (**A**) Schematic representation of the behavior experiment. Mice were injected IP with 4mg prinomastat dissolved in DMSO: 12 hours before fear conditioning, 2 hours after fear induction, and 2 hours after the memory recall test (24h test). In most experiments, memory test was performed 4 weeks later. Each group consisted on 12 mice of mixed gender, and each experiment was repeated 3 times. (**B**1) IP injection of the broad-spectrum MMP inhibitor, prinomastat did not affect fear-conditioning acquisition, tested 24 hours after induction in either FVB WP or FVB MMP9 KO. (**B2**) Memory test 4 weeks later revealed a highly significant difference between control group injected with the vehicle (DMSO) and the experimental groups. MMP9 KO also demonstrated significantly lower freezing percentage in the 4 weeks’ test compared to the WT. MMP9 KO did not exhibit difference between the DMSO and Prinomastat injected groups in the remote-memory test. (**C**) Testing the dynamics of prinomastat on memory indicates an increasing impairment with time. (**D**) Prinomastat impaired remote-memory in C57Bl/6J mice in both cued and contextual tests.

Finally, we tested whether MMPs’ inhibition is specific to FVB or would affect other mice strains. We tested C57Bl/6J that are known to have a better learning ability compared to FVB(27). Our findings with C57Bl/6J were similar to FVB mice.For example prinomastat did not affect acquisition of fear induction as demonstrated in the 24 hours test (Fig S2A). However there was a significant effect on both cue and context tests 4 weeks after conditioning (Fig 4B4). FVB demonstrated minimal contextual memory even 24 hours after fear induction (Supplemental Fig. 2B&C), while C57Bl/6J demonstrated greater memory in both cue and contextual tests. Overall, we observed that PNN proteolysis inhibition did not affect acquisition/recent-memory recall (24hr) but greatly lowered remote-memory (4 weeks).

The temporary inhibition of MMPs with prinomastat followed the schedule and dosage that is described in figure 4A, maintaining sufficient MMPs inhibition to prevent memory consolidation/stabilization. An alternative injection schedule of omitting one or two of any of the injections or lower prinomastat dosage (2mg) were less or not effective. However, ahigher dosage (8mg or 16mg) did not change the degree of memory deficiency (data not shown). Therefore, our data might be compatible with memory stabilization onto PNN-modulation-dependent storage.

The 2h post-retrieval prinomastat injection was required for remote memory impairment, which suggested that MMPs are essential for post-retrieval consolidation into the brain area of the lasting memory location. However, when a single prinomastat dose was injected 2h post-retrieval, there was no remote memory impairment (data not shown).

Weekly memory test after prinomastat injection, demonstrated gradual loss of remote memory (fig. 4C).These results support the hypothesis that PNN erosion is necessary for remote-memory stabilization.

## Discussion

The conservation of life-long memories is essential to sustain vital physiological functions, as well as retain important information acquired as animals experience their environment. Once established, it requires the stability of synaptic input within circuits over very long time. Given that nearly all intracellular proteins turn over rapidly (20, 21), including prominent candidates for memory substrates, we reasoned that a much more stable extracellular structure, such as the PNN,F could play a crucial role in defining, stabilizing, and protecting the synaptic territory. Therefore, PNN could potentially function as a structure that holds the *engram*.

To test the PNN as a stabilizer hypothesis, we characterized the relationship between PNN structure and synapses by EM and 3D SBEM, and demonstrated that PNN holes are almost exclusively occupied by synapses. We applied a SILAM method to test PNN turnover, and established that PNN proteins are long-lasting, while the majority of synaptic proteins are short-live. Also, mice that experienced enriched environment that started at the end of the ^15^N pulse, eroded their PNN faster than their littermates housed in conventional cages, possibly due to learning-induced enlargement of synaptic size and intensity(23).

Additionally, we determined that by inhibition of proteases that erode the PNN, or by using MMP9 knock out mice, there was a significant deficiency in fear conditioning remote-memory, despite normal acquisition. These results support our hypothesis that the PNN could serve to stabilize the code underlying remote-memory information. It also establishes a foundation for further investigation of the PNN in processes involved in age and neurological disease memory impairment.

The role of the PNN in synapse stabilization and memory preservation is still poorly understood despite extensive study over the past century. Camillo Golgi and Ramon y Cajal used several reagents to label a pericellular structure around neurons. Theyspeculated that the structure might be involved in insulating neurons(28). Later studies described PNN composition and the development of new antibodies for Chondroitin sulfate proteoglycans (CSPGs) (29, 30), which were used to investigate individual CSPGs, particularly in the context of glial scar formation(31). The use of ChABC to disrupt PNN structure by degrading GAGs and thus removing the PNN that has been deposited around neurons, resulted in memory loss and reopening of critical periods. This suggests that PNN may regulate experience-dependent synaptic plasticity, maintain remote memories(1, 4, 5, 32, 33) , and play a role in stabilizing drug addiction(34). Abnormal PNN are also correlated with loss or altered memory. PNN are disrupted in brain regions affected by Alzheimer’s disease, and intact PNNs protect neurons when they are exposed to toxic β-amyloid deposits(35, 36). Brain tissue samples from schizophrenia and seizure patients show a reduction of PNNs in affected regions(37, 38). These studies demonstrate that structurally intact PNN is vital to normal brain function, and hints at the possibility that the pattern of holes in the PNN conserves the code for remote-memory(8). This could help us to better understand how PNN changes play a role in normal and pathological brain conditions.

Therefore, we suggest that memories can be preserved in our brains for a lifetime by protecting, defining territories, and stabilize synapses. This argument is supported by observations and experiments, including memory erasure following enzymatic degradation of the PNN using chondroitinase(4, 5, 32), establishment of PNN structure at the closure of critical periods(39, 40), and correlating infantile amnesia to the end of PNN deposition at the closure of critical periods(41–43).

Pyramidal neurons in the hippocampus lack PNN and can be easily modulated(14). However, pyramidal neuronsin the hippocampal CA2 region, are heavily engulfed in PNN, and resist plastisity(44).

An important element of PNN is its stability once it has been deposited, and its resistance to proteases other than MMPs and ADAMTSs(ref). It has also been demonstrated that brevican neoepitopes that result from enzymatic degradation can be found at synapses, and coincide with an increase in synaptic size and strength(45). This indicates that PNN erosion is associated with synaptic strengthening and larger synaptic territory. Our hypothesis suggests that the PNN pattern of holes is a better substrate for stabilizing the *engram.* This is supported by our SILAM studies exploring PNN longevity(20) compared to the shorter-lived intracellular synaptic protein (median turnover rate of less than 4 days).

We found that some of the PNN proteoglycans have the same ^15^N retention as known stable proteins, like histone H4 and MBP. Several synaptic proteins have been implicated in memory maintenance, including PKMζ, CPEB, CaMKII, and Arc/Arg2.1but these proteins have only been examined in relatively recent-memory models and turn over on the time scale of hours to days suggesting a regulatory role rather than a long-term structural one(2, 17, 21, 46, 47).

Intracellular proteins are also susceptible to cellular or brain-wide disruptions that do not necessarily affect memory; including seizure, barbiturate overdose, or metabolic stress, suggesting that they are unreliable substrates for encoding the *engram*. Extracellular proteins remain sequestered away from intracellular ubiquitin and proteasome-mediated degradation and are thought to be less susceptible to disruption.

The mice that experienced enriched environment (EE) created a lasting memory of new experiences; including auditory, tactile, motor skills, social, and olfactory stimulation. The creation of these experiences and memories might have eroded PNN in many parts of the brain, providing a bigger territory for the potentiated and presumably larger/stronger synaptic connections. This may explain the loss of long-lived PNN/ECM components (less ^15^N retention), compared to their littermates that lived in conventional cages (CC)(Fig. 3B, light vs. dark bars). In the EE experiment, we promoted PNN erosion, while in the behavioral experiment we prevented PNN editing and modulation by blocking MMPs activity during learning and consolidation/stabilization.

The results of the SILAM labeling confirmed that the PNN/ECM contains proteins that are extremely stable, as expected from a component that might stabilize the *engram.* The potential removal of ^15^N labeled proteoglycans, subsequently replaced by newly synthetized molecules, could produce enlarged PNN holes. Confirmation of this hypothesis requires microscopic visualization of changes in PNN hole-size in an area that undergoes learning process. or a double-pulsed SILAM experiment using two different stable isotopes.

Varieties of proteases conduct activity-dependent PNN erosion. The proteases required for PNN remodeling have been extensively implicated in synaptic plasticity and LTP. Several groups (see review(48)) have demonstrated that MMP inhibition using antisense oligonucleotides, broad-spectrum pharmacological inhibitors, and neutralizing antibodies can prevent late-phase LTP in various brain regions. Intracerebroventricular injection of a broad-spectrum MMP inhibitor has been reported to attenuate water maze learning for many days after injection. Furthermore, it has been reported that PNN plays a role in regulating fear memory plasticity in the amygdala and visual cortex(1). PNN formation increases in the amygdala between P16 and P45, an interval that corresponds to the establishment of fear memories that are resistant to extinction, and are therefore more persistent(5, 13). In an experiment, that is a reverse of our MMPs inhibition, Gogolla and co-workers(5) removed the PNN by locally injecting ChABC into the basolateral amygdala (BLA) that enabled the extinction of fear memory resembling the extinction effect on young animals at P16 prior to the end of PNN deposition in the amygdala(4).

Here we report that the broad-spectrum MMPs inhibitor, prinomastat, can impair long-lasting memory stabilization. This action could be mainly mediated through the inhibition of MMP9 because MMP9 knockout mice had significantly lower ability to recall fear conditioning 4 weeks after induction yet no further reduction in the memory was induced by prinomastat (fig 4B2).

The temporary inhibition of MMPs with prinomastat might be compatible with memory stabilization onto PNN-modulation-dependent storage. The requirement for a 2h post-retrieval prinomastat injection for remote memory impairment might suggest that MMPs are required for post-retrieval consolidation into the brain area of the lasting memory location which might start at the time that the memory is acquired. Possibly, memories are rapidly encoded into dynamic intrasynaptic molecules that are gradually diminished unless they have been consolidated into stable and permanent storage by forming or enlarging holes in the PNN. These shorter-term memories did not require prinomastat- sensitive MMPs activity. Experiments in human subjects by Bach et al.(49, 50) also demonstrate that MMP inhibition reduced associative memory. Bach et al. blocked MMP-9 with doxycycline showed that MMP-9 is required for structural synapse remodeling involved in memory consolidation. They suggested the use of this treatment as a therapeutic target for unwanted aversive memories since the MMP-9 inhibition attenuated human threat conditioning.

Our results propose some answers to the enigma of *engram* stability and memory conservation. By providing a protected “real-estate,” the PNN holes location and size might provide the code for life-long memories.

## Materials and Methods

Reagents. Primary antibody used rabbit polyclonal anti-Aggrecan (Millipore, Darmstadt, Germany), HRP-conjugated mouse secondary antibody (Cell Signalling, Danvers, MA) was used at 0.4 μg/ml. Fluorescein-conjugated Wisteria floribunda agglutinin (WFA), WFA-biotin, and (Vector Labs, Burlingame, CA) were used at 10 μg/ml. Streptavidin – HRP (VECTASTAIN® Elite® ABC-HRP)

### Electron Microscopy

Mice were anesthetized and perfused with normal Ringer’s solution at 35°C followed by 0.15M cacodylate buffer containing 4% formaldehyde (Electron Microscopy Sciences, Hartfield, PA). Brain tissue was cut into 100 μm vibratome sections. The tissue slices were placed in cold cryoprotectant (78% 0.01M PBS, 20% DMSO, 2% glycerol) for 10 minutes and then they were rapidly plunged into liquid nitrogen for several seconds (until they turned white). Then they were re-immersed in cryoprotectant and allowed to thaw. This cycle was repeated 3 times with the last cycle thawed in cacodylate buffer. We blocked the tissue natural avidin and biotin following the Vector Laboratory protocol. The tissue than was labeled with anti-aggrecan antibody (Millipore) or WFA-Biotin (Vector Laboratories) (20 μg/ml in cacodylate buffer) overnight on a slow shaker at 4°C followed by 3 x 5 minutes wash. The primary Ab was followed by a secondary goat anti-rabbit Ab conjugated to HRP (BioRad) or the WFA-biotin was reacted with streptavidin-HRP (Vector Laboratories). This was followed by 2 minutes incubation of the slices in 0.15M cacodylate buffer containing 2.5% glutaraldehyde (Polysciences). rinsed 5 × 2 min in chilled buffer, then treated for 5 min in buffer containing 20 mM glycine to quench unreacted glutaraldehyde, followed by 5 × 2 min rinses in chilled 0.15M cacodylate buffer. The HRP reaction consisted of a freshly-diluted solution of 0.5 mg/ml (1.4 mM) 3,3′-diaminobenzidine (DAB) tetrahydrochloride or the DAB free base (Sigma) dissolved in HCl were combined with 0.03% (v/v) (10 mM) H_2_O_2_ in chilled 0.15M cacodylate buffer, and the solution was added to cells for 1 to 15 min, depending on the sample(51). To end the reaction, the DAB solution was removed, and cells were rinsed 5 × 2 min with chilled 0.15M cacodylate buffer. For transmitted EM the slices were incubated in 2% osmium tetroxide (Electron Microscopy Sciences) for 30 min in chilled buffer and then rinsed 5 × 2 min in chilled distilled water. Slices were incubated overnight at 4°C in 2% aqueous uranyl acetate (Electron Microscopy Sciences). The slices were then dehydrated in a cold graded ethanol series (20%, 50%, 70%, 90%, 100%, 100%) 2 min each, rinsed once in room temperature anhydrous ethanol to avoid condensation. Durcupan ACM resin (Electron Microscopy Sciences) infiltration wasdone using 1:1 (v/v) anhydrous ethanol and resin for 30 min, then 100% resin 2 × 1 h, then into fresh resin and polymerized in a vacuum oven at 60 °C for 48 h. Areas of interest were identified by transmitted light, and were sawed out and mounted on dummy acrylic blocks with cyanoacrylic adhesive (Krazy Glue, Elmer’s Products). The block trimmed, and ultrathin (80 nm thick) sections were cut using an ultramicrotome (Leica Ultracut UTC6). The sections were imaged on a JEOL 1200 TEM operating at 80 keV.

For Serial block face scanning EM the following SBEM protocol was performed.

### SBEM

Mice were anesthetized and perfused with normal Ringer’s solution containing xylocaine (0.2mg/ml) and heparin (20U/ml) for 2 min at 35°C followed by 0.15M cacodylate buffer containing 2.5% glutaraldehyde (Polysciences), 2% formaldehyde (Fisher Scientific) with 2mM CaCl_2_ at 35°C for 5 min. Brains were removed and postfixed for 18 h at 4°C in the same solution. Brain tissue was cut into 100μm thick sections using a vibratome (Ted Pella) in ice-cold 0.15M cacodylate buffer containing 2mM CaCl_2_, then washed for 30 min in the same solution.

Tissue was placed in a solution containing 1.5% potassium ferrocyanide (Electron Microscopy Sciences) in 0.15M cacodylate buffer with 2mM CaCl_2_ 2% and aqueous osmium tetroxide for 1h at room temperature (RT). Tissue was processed by placing it in 1% filtered thiocarbohydrazide (Electron Microscopy Sciences) for 20 min at R. Then 2% osmium for 30 min at RT, and 1% uranyl acetate overnight at 4°C following triple rinses in DDH_2_O at RT for 30 min. Tissue was triple rinsed in DDH_2_O for 5 min each between each step. We prepared lead aspartate solution (0.066 g lead nitrate; Electron Microscopy Sciences) by dissolved in 10 ml of 0.003M aspartic acid solution, pH adjusted to 5.5 with 1N KOH, warmed in a 60°C oven for 30 min, and filtered. Sections were placed into filtered lead aspartate solution in the 60°C oven for 30 min. The tissue was rinsed five times for 3 min in DDH_2_O and then dehydrated through graded alcohols into acetone and flat-embedded in Durcopan resin (Electron Microscopy Sciences) between mylar strips and placed in a 60°C oven for 48h.

Resin-embedded tissue was mounted on aluminum specimen pins (Gatan) using cyanoacrylic glue and precision trimmed with a glass knife to a rectangle ∼0.5 × 0.75mm so that tissue was exposed on all four sides. Silver paint (Ted Pella) was used to electrically ground the edges of the tissue block to the aluminum pin. The entire specimen was then sputter coated with a thin layer of gold/palladium to enhance conductivity. After the block was faced with a 3View ultramicrotome unit (Gatan) to remove the top layer of gold/palladium, the tissue morphology became visible by back-scattered electron detector imaging and the remaining coating on the edges of the block served to reduce charging. A low-magnification image (∼500×) was collected to identify neurons engulfed with PNN for serial image collection. Tissue blocks were scanned using a Quanta field emission gun scanning electron microscope (FEI) or a Merlin scanning electron microscope (Zeiss) and sectioned at a thickness of 70 or 40nm.

### Image reconstruction

The 3D reconstruction of the images obtained with the SFBSEM were done in two modes; a manual tracing of membranes and organelles in each slice and a semi-automatic segmentation that is described in(52). We used iMod software and the final image was reconstructed using Amira software.

### Stable Isotope Labeling in Mammals (SILAM)

Four FVB females were fed ^15^N based spirulina chow (Cambridge Isotope Laboratories, Inc.) starting at P45 for ten weeks. A male was introduced for one week for mating and the females were kept on ^15^N chow diet during gestation and lactation. At P21, the pups were weaned and were kept on ^15^N diet up to P45 and then switched to^14^N normal chow. The pulse-chase was terminated at different times (months: 0=P45, 6 months=P225, 18 months=P545). For each data point, the brain of a male and female was removed quickly into ice-cold PBS. The brains were cut in half, weighed and half was processed for synaptosomes enrichment and the other half for PNN enrichment. For the six months, pulse-chase a group of 4 P45 pups was transferred into a large cage containing toys, treadmill wheel, mirrors, frequent handling, and extra tactile, auditory, and olfactory stimuli to enhance and enrich their experience.

#### Synaptosomes enrichment

Ten volumes w/v of 0.32M sucrose (Sigma-Aldrich, St. Louis, MO, USA), 4mM HEPES (Sigma-Aldrich) buffer (pH57.4), and cocktail inhibitor tablets (Roche Diagnostics, IN, USA) were added to the brain tissue prior to homogenization with a Teflon pistol attached to a tissue homogenizer rotating at 900 rpm. The tube was slowly moved (up/down) 10 – 15 times until the solution appears homogeneous. The homogenate was centrifuged twice at 1000g for 10 min to pellet the nuclei and the two supernatants were combined and centrifuged at 10,000g for 15 min. The pellet was suspended and centrifuged again at 10,000g for 15 min. This pallet was used as the synaptosome-enriched fraction.

#### PNN enrichment

Brains were homogenized on ice in a glass homogenizer tube in solubilization buffer (10 mM TrisHCl pH 7.5, 150 mM NaCl, 0.5% Chaps, protease inhibitor cocktail (complete, Mini, EDTA-free Protease Inhibitor Cocktail Tablets, Roche) 1tablet/50ml) using a Teflon pestle, at 900 RPM, 10-15 slow strokes, until the tissue looks homogenized. Then the tissue homogenate was sonicate (Sonic Dismembrator 500) on ice (30 sec at 30%) four times. Then the homogenate was centrifuged at 3,900g for 30min in a Sorvall SLA-1500 rotor. The supernatant was then ultracentrifuged at 28,000g for 30 min using a Sorvall SLA-1500 rotor.

The resulting supernatant was loaded on a WFA lectin chromatography (Vector Lab CA, USA) according to Vector lab protocol. The eluate containing the enriched PNN was concentrated by centrifugation in an Amicon Ultra 4, 10K cutoff for 15 min in a swing bucket at 4500 RPM in a Backman counter top centrefuge (Alegra X-22R rotor). This protocol was adapted and modified from Pacharra S. et al (53)

### MudPIT and LTQ Velos Orbitrap MS

Proteomic of PNN and synaptosomes enriched fractions from brains of ^15^N pulse chase experiment are described in details in Butko *et al* (2013) (54).

### Long-Lived protein identification

Described in details in Toyama et al.(22). In short, we identify long-lived proteins in brain fractions (PNN or synaptosomes enriched fractions) by using average peptide enrichments (APE) and a python script to filter the data. Peptides with profile scores < 0.8 were eliminated. Calculation of ^15^N fractional abundance had to be greater than 2.5% to classify a peptide as “m.” All peptides with profile scores > 0.8, but with < 2.5% ^15^N were considered to be “light”. Proteins were considered long-lived if they had more than two “heavy” peptides, more than 65% of the peptides for that protein were “heavy,” and the average ^15^N for all peptides from that protein was > 2.5%. Proteins with a single ^15^N-enriched peptide were discarded as probable misassignments

### Fear conditioning

Littermates FVB wild-type, MMP-9 knockout (FVB background), or C57Bl/6J mice were purchased from Jackson Laboratory and bred at the UCSD vivarium. Each experimental group contained 12 mice of mixed gender and each experiment was repeated 3 times. Fear conditioning was induced at 10 weeks of age using Med Associate Inc. equipment. Mice were introduced into the chamber for 3 minutes of acclamation to the new environment (pre-test freezing evaluation was done during this time). This was followed by a white light and tone (80 dB) for 30 seconds with a scrambled foot shock of 0.8mA at the last second. The light, tone, and shock was repeated 3 times with 1-minute interval. The mice were removed from the chamber 1 minute after the last shock. Mice were injected with 20μl of either vehicle (DMSO) or 4mg of prinomastat in 20μl DMSO 12 hours prior to fear conditioning, 2 hours after fear conditioning, and 2 hours after the 24 hours retrieval test. Retrieval testing was done 24 hours and 4 weeks post conditioning. For the freezing evaluation, we used a video analysis software from Med Associate Inc. with freezing parameters of six frames and threshold motion of 30. Data analysis was done by combining the results from the 3 separate experiments for each condition.

Freezing average, standard error, and p values (2-tailed t-test) were done using Excel program. The data was then Animals that were involved in fights (mostly males) after induction or demonstrated health issues (ear infection) were excluded from the retrieval testing and were not included in the statistical analysis.

## ACKNOWLEDGEMENTS

In memory of Roger Y. Tsien (RYT) who initiated and supported this work. RYT passed away in August 2016. We thank A. Thor and M. Mackay for assistance with electron microscopy sample preparation, and K. Taliman and J. Elbaz for help with behavior experiments. Thanks to Stephen R. Adams for editing the manuscript. This work was supported by NIH grants to RYT, MHE, SRA and VL-R (NS027177), MHE (P41 GM103412) for support of the NCMIR, JY (R01 MH-067880 and P41 GM 103533) JNS (F32AG039127) and (R00DC013805).

## AUTHOR CONTRIBUTIONS

V.L-R, performed the experiments and data analysis. V.L-R., S.P.L. drafted the manuscript. T.J.D, E.A.B, Electron microscopy. J.N.S., D.B.M. mass spectroscopy. A.P., D.R.P., B.H.T data analysis. J.Y., M.H.E., R.Y.T. financial support, editing and guidance. S.S.T. guidelines and editing for final submission. All authors edited the manuscript. R.Y.T. was involved in drafting the paper, however, due to the untimely death he did not have an opportunity to edit the final submitted version.

## SUPPLEMENTARY

**Figure S1.**
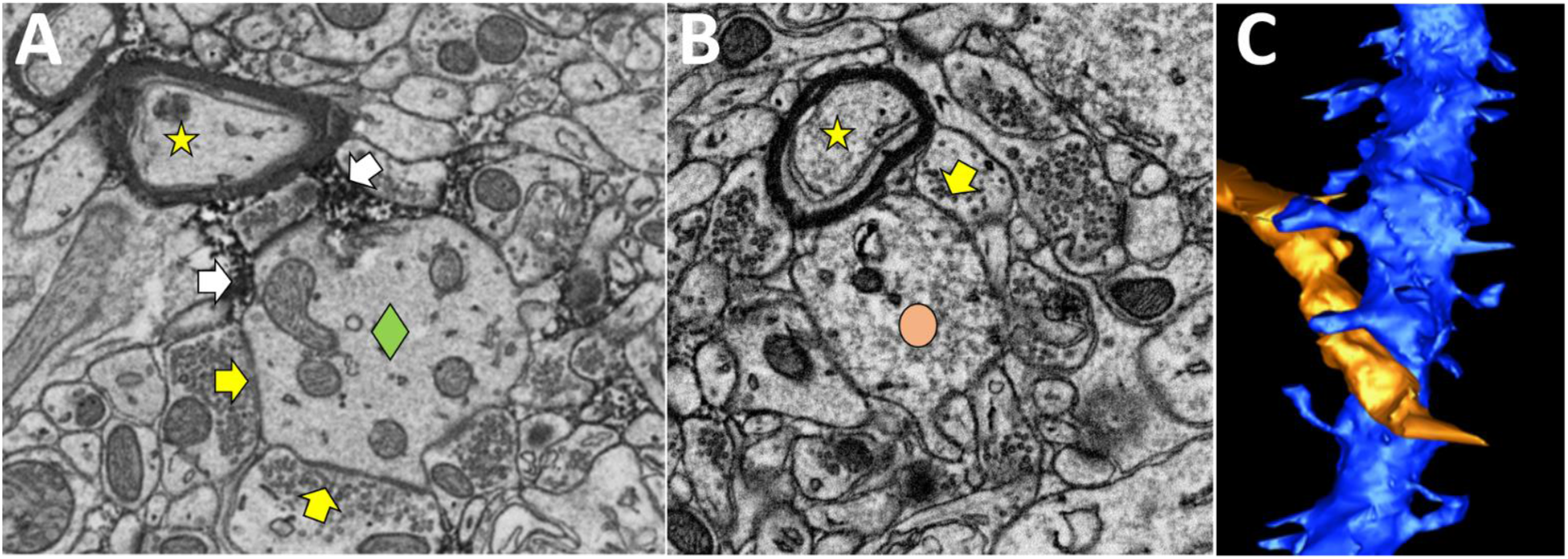
PNN engulfed dendrite surrounded with PNN and synapses. Myelinated fibers that pass by do not touch the dendrite. (**A**) The plasma membrane of a PNN engulfed dendrite () contact, almost exclusively, only pre- synaptic boutons () or PNN (). (**B**) A “naked” spiny dendrite () contacts other cellular elements like a myelinated fiber () and passing dendrites. (**C**) Demonstration of a 3D reconstructed “naked” spiny dendrite (blue) and a passing dendrite (golden) with membrane-to-membrane contact.

**Figure S2.**
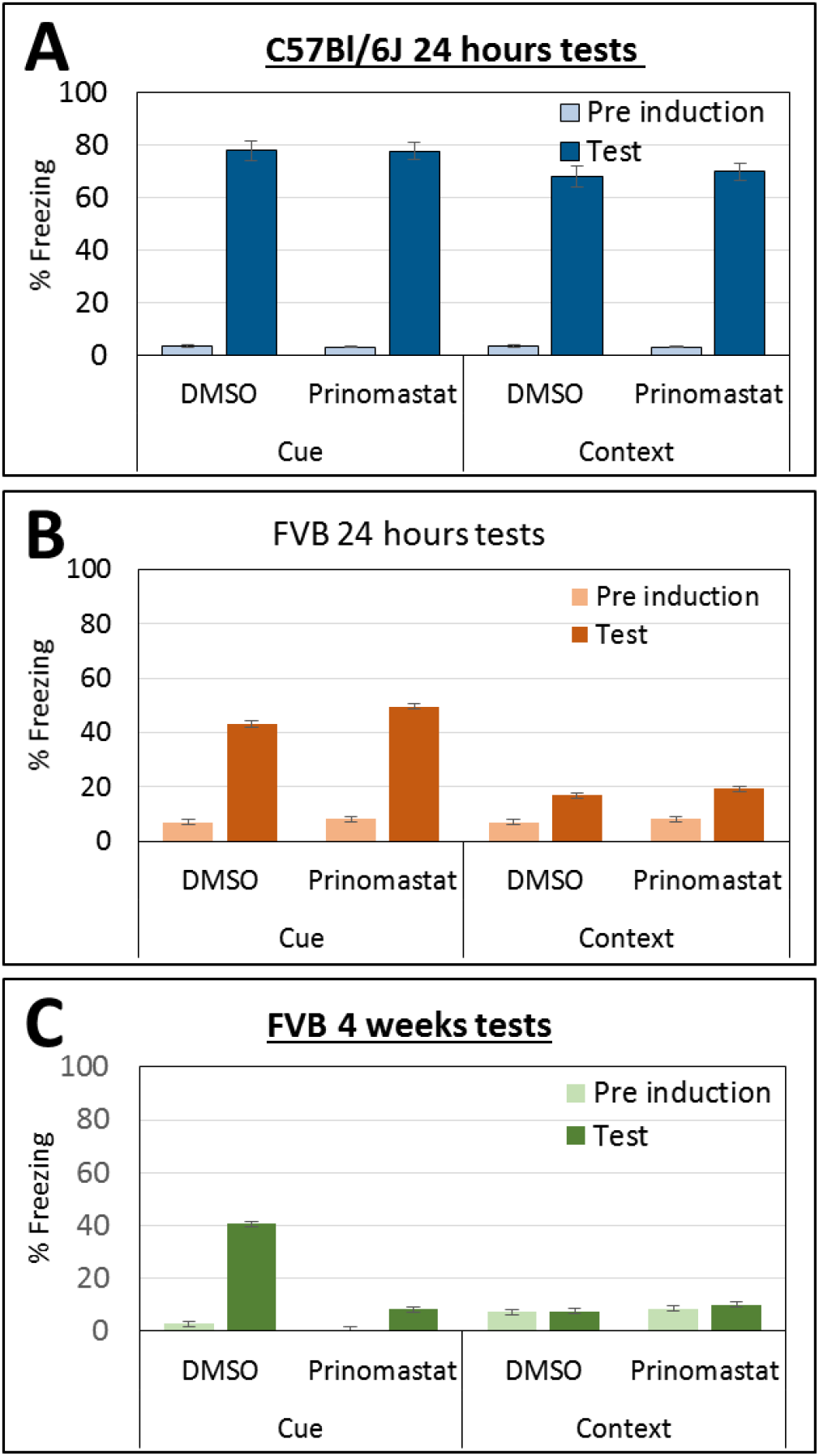
Learning and memory comparison between FVB and C57Bl/6J mice. Fear conditioning memory tested 24 hours after induction demonstrates a higher percentage of freezing by C57Bl/6J mice compared to FVB strain (A&B). FVB mice have very little contextual memory; even 24 hours after fear induction (B), and no contextual memory 4 weeks post induction, while C57Bl/6J mice acquire (S2A) and retain (Fig. 3D) contextual memory.

**Figure S3.**
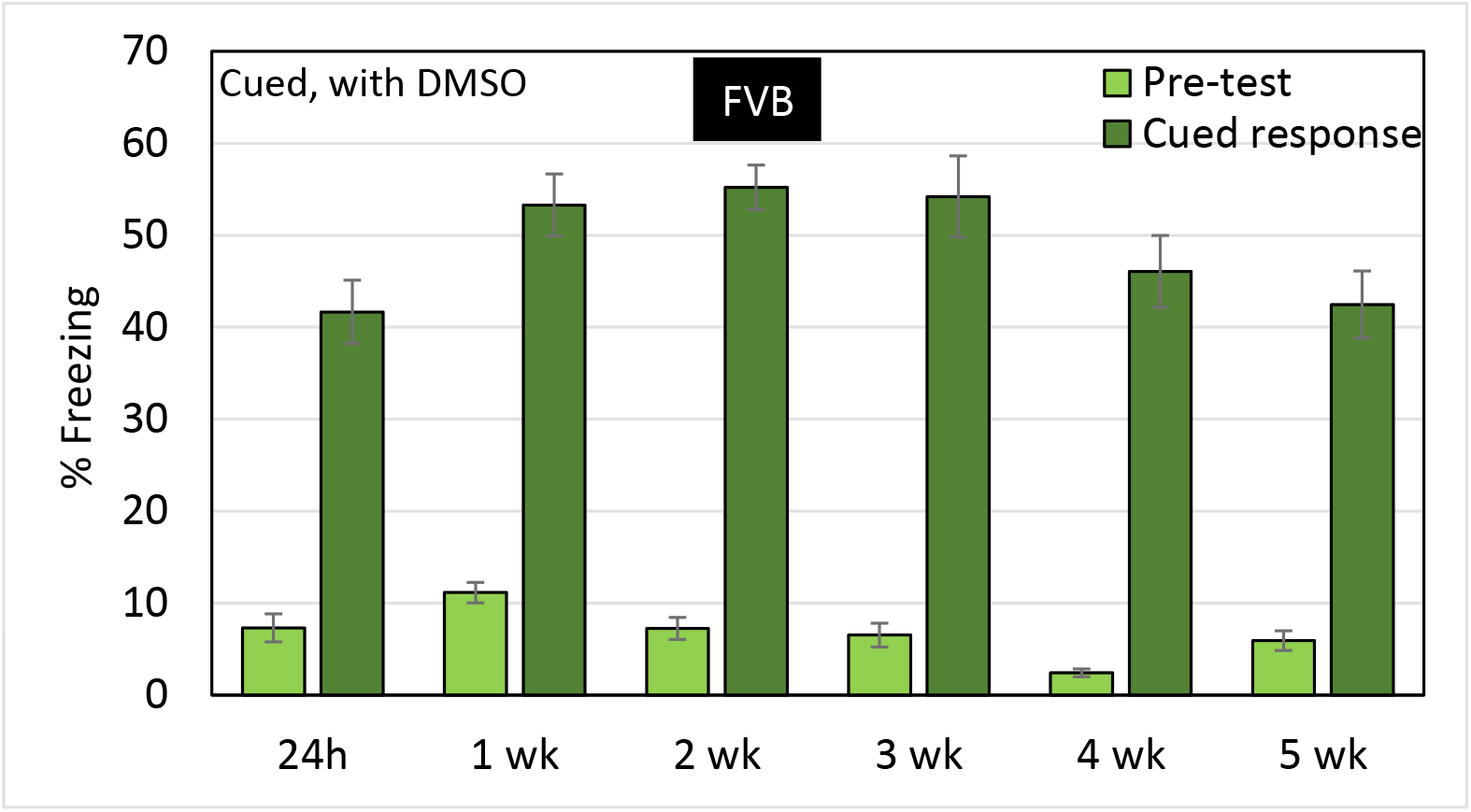
Control groups for the effect of DMSO on memory retention dynamics. In parallel to the prinomastat injection, control cohorts were injected with DMSO at the same time. For each time point, a different group of mice was used to prevent possible extinction of the fear conditioning. These mice demonstrate the persistence of memory for fear conditioning cue test compared to the prinomastat-injected mice (Figure 4C)

**Table S1.** The number of peptides (n) that were identified by mass spectroscopy with ^15^N/^14^N ratio >0.01 that were included in the bar graph of Figure 3.

|  | Conventional cages (6 month pulse -chase) | Enriched environment (6 month pulse -chase) | Conventional cages (18 month pulse -chase) |
| --- | --- | --- | --- |
| CamKII $\alpha,\beta,\gamma,\delta$ | 28 | 40 | 11 |
| Synapsin 1 | 38 | 25 | 21 |
| Synapsin 2 | 27 | 22 | 5 |
| SynCAM 1,2,3,4 | 48 | 28 | 125 |
| CPEB 2,3,4 | 6 | 0 | 7 |
| Aggrecan | 11 | 25 | 48 |
| Brevican | 57 | 50 | 49 |
| Hapln1 | 32 | 46 | 55 |
| Versican | 46 | 32 | 75 |
| Tenascin-R | 99 | 104 | 95 |
| Histone 4 | 40 | 24 | 13 |
| Myelin Basic Protein | 27 | 22 | 94 |

